# Glycogen phosphorylase inhibition alongside taxol chemotherapy synergistically elicits ferroptotic cell death in clear cell ovarian and kidney cancers

**DOI:** 10.1101/2023.05.01.538916

**Authors:** Tashbib Khan, Thomas Kryza, Yaowu He, Jennifer H Gunter, Madeline Gough, Cameron Snell, John D. Hooper

## Abstract

**Background:** Clear cell carcinomas (CCCs) are a distinct histopathological subtype defined by a clear cytoplasm comprised of glycogen and lipids and characterised by poor prognosis and widespread chemoresistance. In the present work we investigate glycogen metabolism as a targetable modality for these cancers.

**Methods and Results:** Adopting the indole carboxamide site pan-glycogen phosphorylase inhibitor CP91149 against clear cell ovarian and renal cancer cell line models, we note antiproliferative and antimigratory effects, as well as energetic stress reflected by reduced ATP pools and increased superoxide-derived reactive oxygen species. Following this, using the agent alongside standard of care chemotherapies for clear cell ovarian (ccOC) and renal cell carcinoma (ccRCC), we note specific synergy with microtubule disrupting chemotherapy paclitaxel, a phenomenon retained in ccOC lines made stably resistant to paclitaxel. Rescue experiments, as well as phenotypic assays suggest that combination-treated cells undergo ferroptotic cell death. We postulate this synergistic efficacy to arise from subjecting the already hypersensitive clear cell cancers to the mitochondrial stress elicited by taxol chemotherapy alongside the oxidative stress augured by glycogen phosphorylase inhibition.

**Conclusions:** Given that CCCs are widely chemoresistant, the present work potentially presents a novel therapeutic avenue for this shared histotype.

## BACKGROUND

Metabolic reprogramming is a hallmark of malignancy (1,2). Tumours adopt unique and varied metabolic phenotypes to adapt to a range of cellular stressors. One such adaptation is the accumulation of glycogen, the stored branched polymer of glucose. Glycogen fuels a range of malignant phenotypes, including proliferation and migration, oxidative stress and senescence protection, histone acetylation, and chemoresistance (3,4). Glycogen metabolism is upregulated in several oncogenic contexts, including hypoxia, via the *RAB25A* oncogene, and in a unique histotype of carcinomas known as the clear cell cancers (CCCs) (5–7).

CCCs are a distinct histopathological subtype defined by a clear cytoplasm comprised of glycogen and lipids and characterised by poor prognosis and widespread chemoresistance (8). The two most common CCCs are clear cell ovarian carcinoma (ccOC), the secondmost common and most chemoresistant ovarian cancer, and clear cell renal cell carcinoma (ccRCC), the most frequent and metastatic kidney cancer, although CCCs have been noted in diverse tissues of origin, including breast (9), colorectum (10), and endometrium (11). Only recently have parallels begun to be drawn between CCCs beyond the conspicuous histology, with studies highlighting common origins in hypoxia, shared metabolic states, and hypersensitivity to glutathione peroxidase 4 (GPX4)-dependant ferroptosis (3,8,12). Although inroads have been made in understanding and targeting the role of lipids in this shared context, glycogen has been given scarce attention. Hence, the present work aims to investigate glycogen as a therapeutic target in CCCs.

In ccOCs, which are thought to originate in the hypoxic and oxidative stress environment of endometriotic cysts, glycogen accumulation arises as a stress-response mechanism via the overactivation of the transcription factor *HNF-1B* (13,14). Stored glycogen facilitates cell growth, cellular detoxification, and platinum resistance (13,14). Recently, we reported that concentrations of the small molecular weight glucose analog 2DG that are too low to disrupt cell proliferation can nevertheless disrupt glycogenolysis when combined with platinum chemotherapy (15). Recent work has also highlighted the importance of glycogen to the cross-talk between ccOC cancer cells and cancer-associated fibroblasts (16–18). In ccRCC, upregulated glycogen metabolism is thought to occur because of VHL deficiency, giving rise to constitutively active HIF1α and HIF2α, resulting in glycogen accumulation (12,19,20). This glycogen metabolism signature is reflected at both the genetic and protein levels of ccRCC (21–24). Recently glycogen synthase (GS) was shown to be an independent predictor of poor prognosis and mediator of resistance to the tyrosine kinase inhibitor sunitinib in *in vitro* and *in vivo* models of ccRCC (25). However, it is important to note that another recent study reported that glycogen metabolism is dispensable for primary tumour growth in ccRCC (26).

The present work examined glycogen metabolism as a therapeutic target in both ccOC and ccRCC. It characterises the importance of glycogen and examines inhibitors of a key enzyme of glycogen catabolism, glycogen phosphorylase (PYG), in ccOC and ccRCC cell line models. Aberrant expression and function of the enzyme has previously been noted in cancer as well as diabetes and other metabolic disorders, rationalising the development of PYG inhibitors (27). These inhibitors have been tested largely in diabetes models but have scarcely been examined in cancer models (27–30). Hence, one of the goals of the present work was to assess whether CCCs, with their unique cytoplasmic glycogen stores, could provide a therapeutic window for PYG inhibition. In this vein, we pursued the PYG inhibitor CP-91149 in cell line models of ccOC and ccRCC, finding dose-dependent energetic stress and apoptosis akin to that observed in previous studies. We also assessed the ability of PYG inhibition to synergise with existing chemotherapeutic modalities of ccOC and ccRCC, and interestingly found robust synergy with microtubule-disrupting chemotherapy paclitaxel, potentially auguring ferroptosis. Given they bear few effective therapies, the combination of paclitaxel and PYG inhibitor can garner a new therapeutic avenue for CCCs.

## METHODS

### Reagents and Antibodies

The following antibodies were from Cell Signalling Technologies: Phospho-Glycogen Synthase (Ser^641^) #3891, total glycogen synthase #3886, Phospho-AKT (Ser^473^) #9271, AKT (pan) #4691, Phospho-GSK-3β (Ser^9^) #9336, GSK-3β #9315, GAPDH #2118, Anti-rabbit IgG (H+L) (DyLight™ 800 4X PEG Conjugate) #5151, Anti-mouse IgG (H+L) (DyLight™ 680 Conjugate) #5470. Antibodies against PYGL (PA5-9152) and PYGB (PA5-28022), dyes for TMRE (T669), Calcein-AM (C3100MP), PI (PP1304MP), BODIPY™ 581/591 C11 (D3861) and MitoSOX-PE (M36008) were from Thermo Fisher Scientific. PE Anti-human TfR1 antibody (#334106) and FITC Annexin V (#640906) were from Biolegend. All antibodies were used at 1:1000 for immunoblotting, except for GAPDH which was used at 1:10000, and Anti-rabbit IgG 800 and Anti-mouse IgG 680 which were used at 1:20000. For flow cytometry and fluorescent microscopy, antibodies were used according to manufacturer’s instructions.

The following inhibitors were obtained from SelleckChem: CP91149 (#S2717), Sorafenib (#S7397), Z-VAD-FMK (#S7023), Necrostatin-1 (#S8037), Deferoxamine mesylate (#S5742). Mitomycin C (#M4287), BAY R3401 (#B3936), BI2596 (#S1109), Nocodazole (#S2775) and Chloroquine diphosphate salt (#C6628) were from Sigma-Aldrich. Carboplatin and paclitaxel were from Mater Pharmacy. All reagents were reconstituted according to manufacturer’s instructions.

Tissue culture plastics and medium, as well as immunoblotting gels and transfer stacks, were from ThermoFisher Scientific. ImageLock plates (96-well; #4379) as well corresponding Woundmaker Incucyte® Cell Migration Kit (#4493) were from Sartorius. All other consumables were from Sigma-Aldrich except where noted.

### Cell Lines and Culture Conditions

As mentioned above, in sections 3.4.2. and 3.6.4.2, 786-O and A498 ccRCC cells were from ATCC (Manassas, VA, USA). OVTOKO cells were kindly provided by Dr Katherine Roby (University of Kansas School of Medicine, Kansas and KOC7C cells were courtesy of Dr. Hiroaki Itamochi (Tottori University School of Medicine, Yonago, Japan). All lines were cultured in RPMI-1640 with 10% heat-inactivated fetal calf serum, and 1X penicillin-streptomycin.

### Luciferase-labelling of cells

Luciferase labelling of cells for in vivo experiments was performed as outlined in Section 3.4.3. Briefly, cells were stably transduced with a lentivirus-based expression construct generated by cloning a polymerase chain reaction (PCR)-amplified DNA fragment encoding luciferase from a pGL4.10-lucIIplasmid (Promega, Sydney, Australia) into a pLenti CMV Hygro DEST vector (Addgene, Cambridge, MA, USA) using Gateway LR recombination cloning technology (Life Technologies, Mulgrave, Australia), as previously described (117). Cells stably transduced with the luciferase construct were selected in puromycin (2 μg/mL; Thermo Scientific).

### Western Blot Analysis

Whole-cell lysates treated under stipulated conditions were collected using RIPA buffer (Sigma-Aldrich) with 1 × Complete protease inhibitor cocktail (Roche, Castle Hill, NSW, Australia), 2 mM sodium vanadate, and 10 mM sodium fluoride, and used in western blot analysis as described previously. Briefly, Lysates (20-40 μg for cells and 80µg for tissues) were separated by SDS-PAGE under reducing conditions (except where noted), transferred onto nitrocellulose membranes, and blocked in fish skin gelatin blocking buffer (Sigma Aldrich) (3% w/v in PBS, with 0.1% Tween-20 and 0.05% sodium azide). Membranes were incubated with primary antibodies diluted in blocking buffer overnight at 4°C, washed with PBS containing 0.1% Tween 20 (Sigma-Aldrich), and then incubated with appropriate secondary antibody. Signals were detected using an Odyssey Imaging System and software (LI-COR Biosciences, Millennium Science, Mulgrave, Australia).

### Endpoint Cell Count Measurement of Cytotoxicity

Endpoint Cell Count Measurement of Cytotoxicity were performed as noted above. Briefly, following specified cell treatments, media was removed and cells were fixed with 50μl ice-cold methanol for 5 minutes. Methanol was then removed, cells were washed in PBS, and then stained with 0.5 μg/mL DAPI (Thermo Scientific) for at least 30 minutes. Cells were then imaged at 4X using the InCell 2200 or 6500 automated fluorescence microscope system (GE Healthcare Life Sciences) and quantitative analysis was performed with Cell Profiler Software(182,183). Cell numbers were normalized to untreated controls and plotted as “relative cell number.”

### Cell Viability Assay

Cell viability was measured as outlined in Chapter 3. Briefly, cells (1500 cells/well) were plated in triplicate in 96 well plates and allowed to attach overnight, before treatment for 72 h with stipulated drug concentrations. Cell viability was assessed using a CellTiter cell proliferation assay kit (Promega), according to the instructions of the manufacturer. Tetrazolium dye solution (20 μL) was added to each well, followed by incubation at 37 °C for 3 h, then measurement of absorbance at 490 nm, using a PHERAstar Omega plate reader (BMG Labtech, Mornington, Australia), as previously described (25,117,220).

### Incucyte measurements of cell proliferation

Cells (3,000 per well) were plated in 96-well plates and allowed to attach overnight. The following day, cells were treated with stipulated drug concentrations and by imaging on the Incucyte Zoom S3 (Sartorius). Images were taken every two hours and fold change in confluency calculated using Incucyte software.

### Wound Healing Assay

Cells (50,000 per well) were plated in 96-well ImageLock plates (Sartorius) and allowed to attach overnight. The following day, cells were pre-incubated for 6 hours with stipulated drug concentrations and for 2 hours with mitomycin C (0.5 μg/mL) to halt proliferation. Cells were then scratched using the Essen Biosciences automated Woundmaker according to manufacturer instructions (at least two scratches per plate). Cells were washed twice in warm PBS and replenished with drug-containing media, following by imaging on the Incucyte Zoom S3 (Sartorius). Images were taken every two hours and relative wound confluence (%) calculated using Incucyte software.

### Colony Formation Assay

The colony-forming ability of cell lines was assessed as previously (117). Briefly, Cells (100,000/well) were seeded overnight in six well plates, followed by treatment for 72 h with indicated drug concentrations. Here, Paclitaxel GI_50_ represents the paclitaxel dose defined by cell count measurements of cytotoxicity (100nM for OVTOKO, 250 nM for KOC7C. 290 nM for 786O and 1 μM for A498). After 72 hours, cells were non-enzymatically dissociated, and then replated at low density in a 24-well plate (500 cells/well), followed by growth for 1–2 weeks. The media was then removed from the plates and washed gently with PBS twice, followed by fixation with 4% paraformaldehyde (Sigma-Aldrich) for 30 minutes. The cells were then washed gently in PBS and stained with 0.1% crystal violet (Sigma-Aldrich). The stain was removed after at least 30 mins and plates were washed under running RO water, allowed to dry overnight, and then scanned.

### Glycogen Content Assay

Total cellular glycogen content was measured as previously (117,179). Briefly, cells treated for stipulated drug doses were scraped in glycogen isolation buffer (Tris pH 8, 150 mM NaCl, 50 mM NaF, 2 mM EDTA, 5 mM sodium pyrophosphate). Samples were then homogenized by passing through 26-G needles and cleared by centrifugation at 14,000 g and 4°C for 30 min. Protein concentration was quantified by micro-bicinchoninic acid assay (Thermo Fisher Scientific), and then glycogen quantification assay was performed as previously. Briefly, 5 μL of amyloglucosidase (Megazyme, 3260 Units/mL) was added to the reaction mixture containing 10 μL of sample, 20 μL of 200 mM sodium acetate buffer (pH 4.5), 2.5 μL of 10% acetic acid and 62.5 μL of deionized water. Two controls with 10 μL of deionized water (instead of homogenate) were also analysed. Samples were incubated for 30 min at 50 °C and then centrifuged at 20 000 g for 20 min, with the supernatant being transferred to a new tube and the pellets being discarded. Then, 5 μL of degraded glycogen and 5 μL of D-glucose standards up to 1 mg/mL was mixed with 170 μL of reaction buffer containing: 150 μL of 200 mM tricine/KOH (pH 8) and 10 mM MgCl_2_; 18 μL of deionized water; 1 μL of 112.5 mM NAPD; 1 μL of 180 mM of adenosine triphosphate (ATP) and 0.5 U of Glucose-6-phosphate dehydrogenase(G6PDH) (Roche). After the absorbance at 340 nm was recorded for 20 min to determine a baseline, 4 μL of Hexokinase solution (0.75 U in 5 μL of 200 mM tricine/KOH (pH 8) and 10 mM MgCl_2_) was added to each well and the absorbance at 340 nm was recorded for 30 min. Glucose concentration in each well was calculated by subtracting the baseline absorbance from the absorbance plateau, using the D-glucose standard curve, and water blank background values were subtracted. This was then used to extrapolate the glycogen concentration in the initial sample. Samples unreacted with amyloglucosidase were used to verify the insignificance of background glucose.

### Bioenergetic Analysis of ATP Production

ECAR and OCR, and specifically ATP derived from glycolysis and mitochondrial oxidative phosphorylation, were quantified in real time using a Seahorse XFe Extracellular Flux Analyser (Seahorse Bioscience, Agilent, In Vitro Technologies, Noble Park North, Australia), as described previously, and above in section 3.4.8(117). Here, cells were treated for 24 h with either 20 μM or 100 μM CP91149. On the day of the assay, the cells were washed twice with XF RPMI media (Seahorse Bioscience), then cultured in this media supplemented with 10 mM glucose, 2 mM glutamine, 5 mM HEPES, and 1 mM sodium pyruvate (pH adjusted to 7.4 + 0.05). Cells were then cultured in a non-CO2 incubator for 1 h prior to the start of the experiment, when rotenone and antimycin A (5 µM each) were injected into the media to eliminate cellular citric acid cycle activity, followed by oligomycin (100 mM) to eliminate all ATP production, with differences enabling the calculation of ATP derived from both glycolysis and mitochondrial oxidative phosphorylation. The assay plates included control blank wells containing only media to which the various reagents were added for experimental wells. Measurements from blanks were automatically subtracted by the instrument Wave 2.6 software. Data were normalized to the cell number per well, which was determined by imaging of DAPI stained cells using the Incell 2200 or 6500 Cell Imaging Systems (GE Healthcare, Parramatta, Australia).

### Flow Cytometry

Cells (100,000 per well) were seeded in 6 well plates and allowed to attach overnight. The following day, cells were treated with stipulated drug concentrations for 24 hours.

For MitoSox Red and calcein-AM assays: On the day of the assay, cells were lifted non-enzymatically and incubated for 30 minutes with either 5μM MitoSoX Red, 100nM calcein-AM in the dark in a tissue culture incubator. Cells were then washed twice in warm PBS and then analysed using a flow cytometer (FACsFortessa, BD Biosciences) in the PE channel for MitoSox Red and the FITC channel for calcein-AM. At least 10,000 events were analysed per condition.

For BODIPY 581/591 C11 assays: On the day of assay, 5μM BODIPY 581/91 C11 was added to wells, and cells were incubated for 45 minutes in a tissue culture incubator in the dark. Then, cells were lifted non-enzymatically, washed twice, and analysed in both a FITC and PE channels using a flow cytometer (FACs Fortessa, BD Biosciences), with at least 10,000 events analysed per condition.

For TfR1 assays: On the day of the assay, cells were lifted non-enzymatically, blocked for 30 minutes in 0.5% bovine serum albumin in PBS at 4°C, and then incubated for one hour in PE anti-human CD71 Antibody in the dark at 4°C (1:100; Biolegend). Cells were washed twice with PBS and then and then analysed using a flow cytometer (FACsFortessa, BD Biosciences), with at least 10,000 events analysed per condition.

### Synergism Measurements

Synergism between drug treatments was measured using the Chou-Talalay Combination Index Method using the Compusyn software (Combosyn Inc.) (https://www.combosyn.com/), as described above. CI values <1 were considered synergistic interactions, 1 additive interactions, and >1 antagonistic interactions.

### Generation of Paclitaxel Resistant Lines

The generation of OVTOKO and KOC7C cells stably resistant to paclitaxel (Pac_R_) was performed based on GI_50_ values calculated for both lines using the above cell count measurement of cytotoxicity (100 nM and 250 nM respectively). Cells were treated in T25 flasks with stepwise doses of paclitaxel starting at 1/10 GI_50_ doses. Cells were treated for at least 3 days, followed by change into drug-free media for at least 4 days to allow resistant cells to re-proliferate. This was repeated for doses ¼ GI_50_, ½ GI_50_ and at GI_50_. Once cells displayed stabilized growth in the presence of GI_50_ doses of paclitaxel they were expanded and assays performed alongside passage-matched parental cells (Pac_Control_). Cells were routinely re-treated with GI_50_ values of paclitaxel to ensure they did not revert to parental sensitivities.

### Time-Lapse Annexin-V/PI Imaging

Time-lapse imaging of cell death was performed as previously describe(198). Briefly, cells (4000 / well) were plated in phenol-red free complete DMEM and allowed to attach overnight. On the day of assay, Cells were preincubated for 30 minutes in annexin V-FITC (1 μg/ml; BD Biosciences) and Propidium Iodide (1 μg/ml; Sigma Aldrich), treated with stipulated drug concentrations, and then imaged every 30 minutes on the Incucyte Zoom S3 (Sartorius). Cell-by-cell analysis was performed specific to every line using Incucyte software, with cells demarcated into the following 4 groups: 1) Green – Red –, 2) Green + Red –, 3) Green – Red +, and 4) Green + Red +.

### Mitochondrial Membrane Potential

Cells (10,000/ well) were seeded in 96 well plates in phenol red-free growth media and allowed to attach overnight. On the day of assay, cells were loaded with TMRE (200 nM) for 30 min at 37 °C before indicated treatments. Cells were then imaged on the Incuyte Zoom S3 at 15-minute intervals using the phase contrast and Red Channel (Excitation 585 nm [565,605]). TMRE fluorescence per cell was calculated using the Incucyte cell-by-cell analysis. When fluorescence changes had plateaued the analysis was stopped, and the steady-state fluorescence/cell /well was plotted (209).

### Rescue Experiments

The capacity of common inhibitors of cell death modalities (zVAD-FMK; apoptosis, necrostatin-1, necroptosis; chloroquine, autophagy; deferoxamine, ferroptosis) to rescue combination efficacy was assessed. Cells were treated as outlined with paclitaxel (0-10 μM) with or without a fixed 20μM dose of CP91149, in the presence or absence of either zVAD-FMK (20μM), necrostatin-1 (20μM) or chloroquine (50μM). Cells were then fixed after 72 hours, stained with DAPI and enumerated as outlined above in section 4.4.5.

### Statistical Analysis

Except where noted, the Mann-Whitney test was used in analysis comparing two groups while the Kruskal-Wallis test was used for comparisons involving more than two groups. A value of p ≤ 0.05 was considered significant. Significance values are represented in graphs as *p < 0.05, **p < 0.01, ***p < 0.001 and ****p < 0.0001. Data and statistical analyses were performed using Prism 6.0 software (GraphPad, San Diego, CA, USA).

## RESULTS

### PYG inhibitor CP91149 impairs growth and elicits energetic and oxidative stress in clear cell cancer lines in vitro

To directly examine the role of PYG in CCC, we disrupted its function pharmacologically using CP91149. This agent inhibits the indole carboxamide site of the enzyme and has previously been used for both cancer and diabetes studies (31–33). Importantly, the inhibitor successfully targets all three PYG isoforms, so circumvents isoform redundancy (28). We first assessed the efficacy of CP91149 in ccRCC lines 786-O and A498 and ccOC lines OVTOKO and KOC7C by treatment for 72 hours with increasing concentrations of the compound, followed by fixation, staining with DAPI, and automated cell counting. Figure 1A shows that the inhibitor elicited a concentration-dependent reduction in cell number for all four lines, with the two ccOC lines displaying a lower GI_50_ (60 μM and 65 μM) than the two renal lines (94 μM and 86 μM). Then, the effect 20 μM or 100 μM of the on the migration and clonogenic potential of the four CCC lines inhibitor was assessed. The two concentrations (highlighted by broken lines in Figure 1A) were selected because 100 μM inhibits the PYG enzyme in cell-free systems (28,31) and exerted over 50% reduction in cell number in Figure 1A, whereas 20 μM was largely ineffective at exerting antiproliferative effects. *In vitro* migration was next assessed by measuring the ability of a confluent cell monolayer to heal a scratch wound. Figures 1B shows that 100 μM CP91149 impinged the wound healing capacity of the two ccOC lines OVTOKO and KOC7C (p<0.05), but that this same dose had minimal effect on ccRCC lines 786-O and A498; possibly a reflection of the higher doses required to exert efficacy for ccRCC. 20 μM CP91149 had little effect on any of the lines. The effect of CP91149 on the clonogenic survival of cells was then assessed. Cells were treated for 72 hours with indicated concentrations, and then re-plated in drug-free media for 7-14 days, and colonies were stained with crystal violet and colony area quantified using ImageJ. Figure 1C shows that 20 μM CP91149 had no impact on colony formation of any of the lines, whereas 100 μM of the inhibitor attenuated clonogenic survival of ccOC lines OVTOKO and KOC7C by 30%, and ccRCC line A498 by 70% (p<0.05). Interestingly, there was no effect of either dose on ccRCC line 786-O. Generally, CP91149 seems more effective at reducing proliferation, migration, and colony formation in ccOC lines than ccRCC lines, consistent with the lower GI_50_ observed.

**Figure 1.**
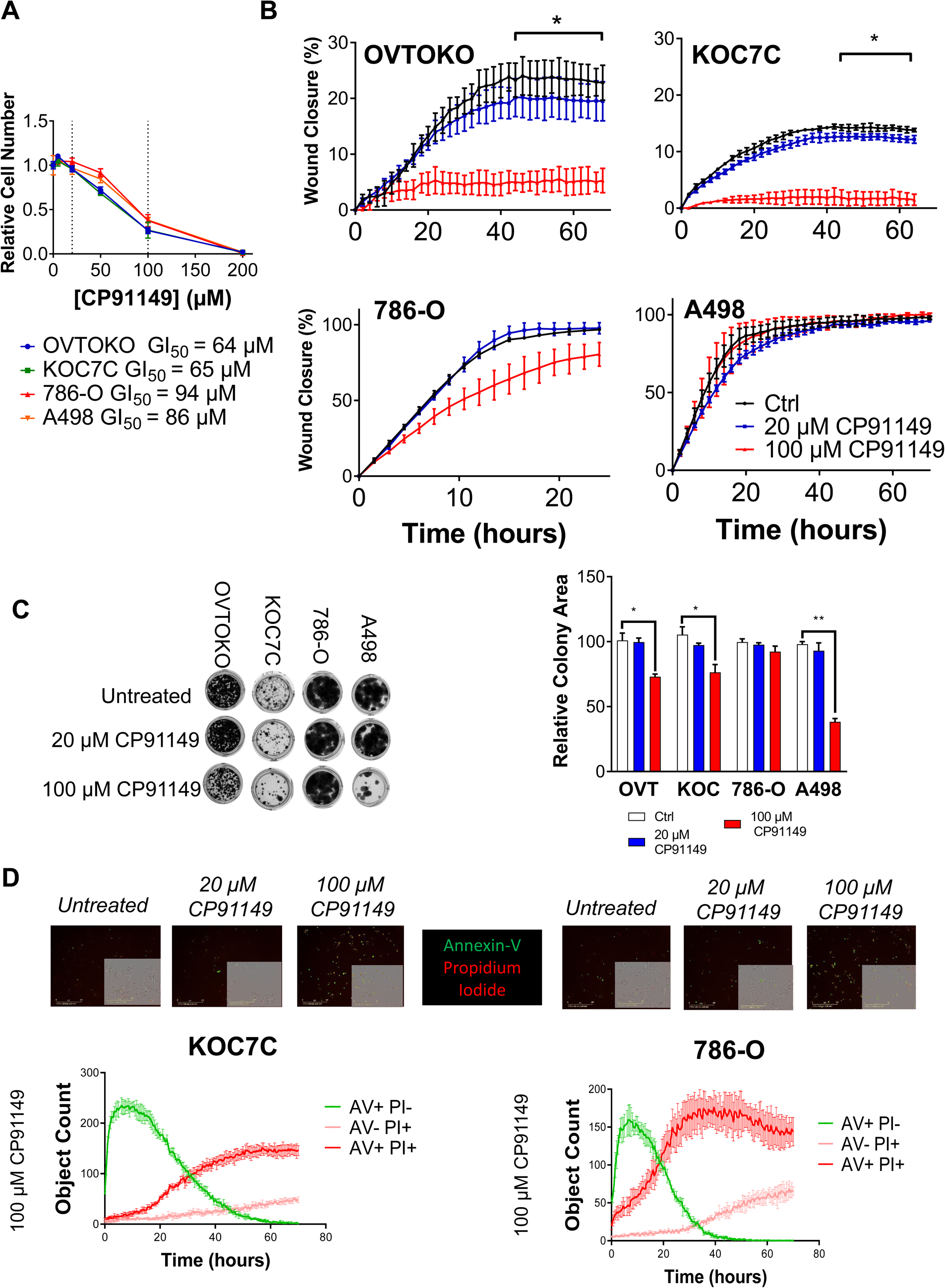
CP91149 attenuates proliferation, migration and clonogenic survival in CCCs. **A)** CP91149 monotherapy efficacy was assessed in clear cell carcinoma lines. 3000 cells/well were plated and allowed to attach overnight. Cells were then treated with indicated concentrations of CP91149 for 72 hours, followed by methanol fixation, DAPI staining and cell counting on the Incell 6500 system and analysis using CellProfiler software. **B)** Cells (50,000/well) were allowed to attach overnight. On the day of the assay, cells were pretreated for 12 hours with relevant drug concentrations, as well as Mitomycin C (10 μg/mL for 2 hours). Cells were then scratched using an Essen Woundmaker, washed, replenished with drug-containing media, and imaged every two hours using the Incucyte S3 system. **C)** Cells (50, 000 per well) were plated in 6 well-plates and allowed to attach overnight. Cells were then treated with relevant drug concentrations for 72 hours, after which time cells were lifted and replated (1000/well) in drug-free media, and grown for 2 weeks, then fixed and stained with crystal violet, imaged, and relative colony area calculated using ImageJ. **D)** Cells (4000/well) were plated in phenol red free DMEM medium and allowed to attach overnight. On the day of assay pre-incubated in phenol red-free DMEM media containing annexin V (AV; 1 μg/ml) and propidium iodide (PI; 1 μg/ml). Cells were then treated with indicated concentrations of CP91149 and imaged every 30 minutes using the Incucyte S3 system. Representative images from 72 hours are shown. Single cells were analysed using Incucyte software and defined as either AV+PI-, AV-PI+, AV+PI+ or AV-PI-, and cell populations are plotted with time. Data is presented as mean ± SEM and was analysed using a Mann-Whitney test with *p<0.05 **p<0.01, ***p<0.005 and ****p<0.001

CP91149 also elicits apoptosis in models of hepatocellular carcinoma (32). To assess if the drug was displaying a similar mode of action in CCCs, KOC7C ccOC and 786-O ccRCC cells were stained with annexin V-FITC and propidium iodide (PI), treated with CP91149, and time-lapse imaged using the Incucyte S3 system. Annexin-V stains membrane-protruded phosphatidylserine, an early event in programmed apoptosis, whereas PI binds the nuclei of cells with perforated plasma membranes; a conserved marker of late-stage cell death (34,35). Thus, apoptosis traditionally entails early-stage annexin-V positivity followed by later annexin-V/PI dual-positivity. Here, the proportion of cells positive for annexin-V (AV) and PI were calculated using Incucyte software, and cells were demarcated into AV+PI, AV-PI+, AV+PI+ or AV-PI-, and these were plotted with time. Figure 1D highlights endpoint representative images of cells treated with the two doses of inhibitor. Below this, the dose-time-dependant effect of 100 μM CP91149 is highlighted, showing that at high doses cells displayed a robust early increase in annexin V positivity followed by dual annexin V/PI positivity, corroborating a time-dependant induction of apoptosis.

### CP91149 increases glycogen levels, reduces ATP levels and generates superoxide ROS

Next, we assessed the effect of CP91149 on total glycogen levels, to corroborate that the inhibitor was exerting an on-target effect. Changes in total cellular glycogen levels were measured for cells treated for 24 hours with 20 μM or 100 μM CP91149. A shorter timepoint was selected for these assays to overcome the cytotoxicity observed for high doses of the inhibitor at 72 hours in Figure 1. Figure 2A shows that the inhibitor causes an increase in glycogen levels for all four lines. This increase is probably reflective of a homeostatic response in glucose flux towards glycogenesis, a phenomenon previously noted by Favaro et al. (2012) with *pygl* silencing under hypoxic conditions (7). Interestingly, the effect more pronounced in the ccRCC lines 786-O and A498, with both 20 μM or 100 μM CP91149 doubling glycogen levels (p<0.05); contrastingly, for ccOC lines OVTOKO and KOC7C, only 100 μM CP91149 showed significantly increased glycogen levels (p<0.05).

**Figure 2.**
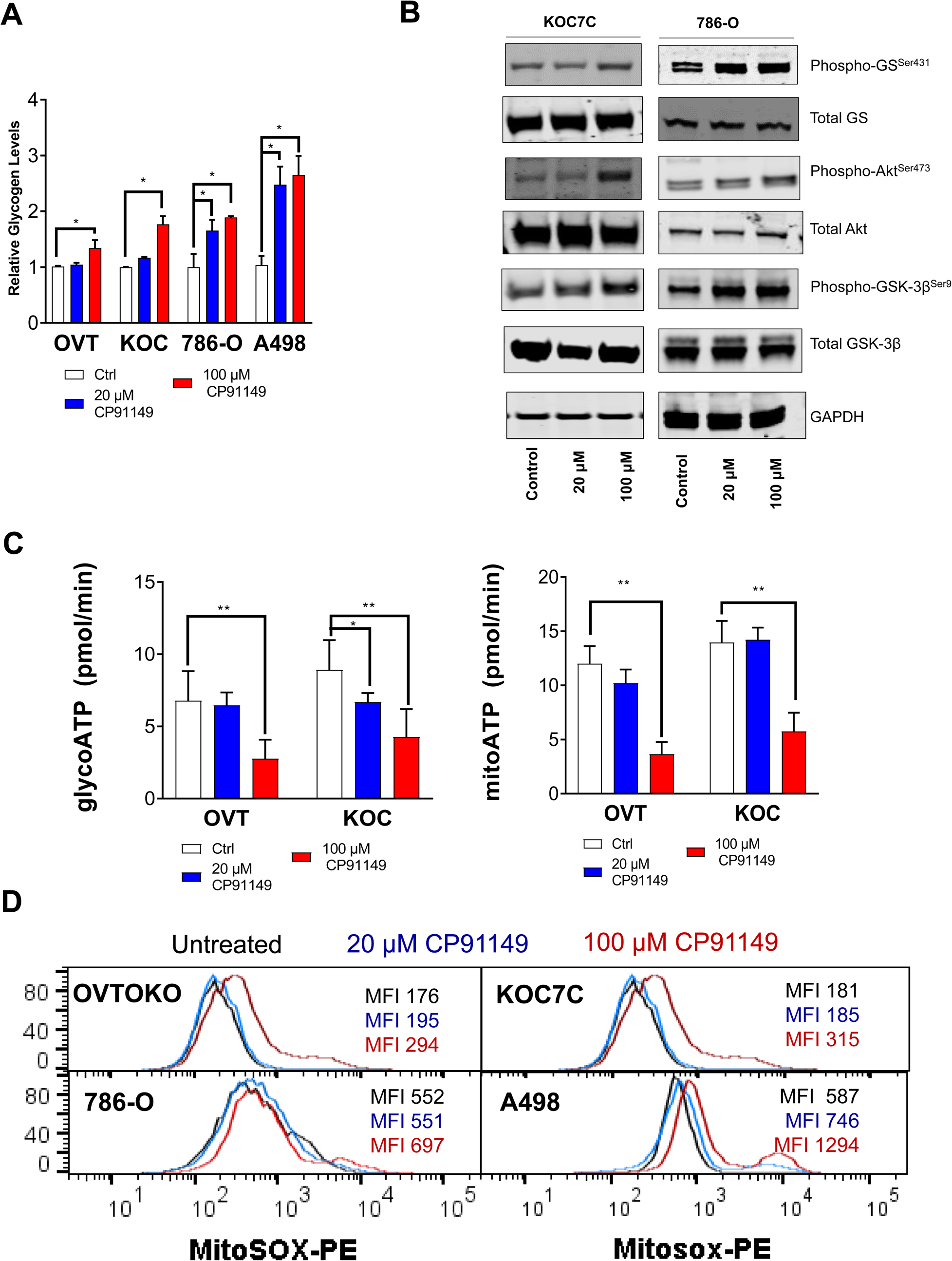
CP91149 increases glycogen levels, reduces ATP levels and generates superoxide ROS. A) OVTOKO (OVT), KOC7C (KOC), 786-O and A498 cells were treated for 24 hours with relevant drug concentrations and assayed for glycogen; Cells were lysed in glycogen isolation buffer, glycogen degraded using amyloglucosidase, and glycogen-derived NADPH detected spectrophotometrically. Glycogen levels were normalized to untreated control. B) Cells were treated overnight with relevant drug concentrations, lysed in RIPA buffer, and immunoblotted for phospho Glycogen Synthase^s641^, total Glycogen synthase, phospho Glycogen Synthase^s641^, total Glycogen synthase phospho-GSK3B^s9^, and total GSK3B. GAPDH was used as a loading control. C) Cells were treated overnight with relevant drug concentrations and the Seahorse ATP Rate assay was performed. Glycolysis-derived ATP (GlycoATP) and oxidative phosphorylation-derived ATP (MitoATP) are shown. Data was compared using a Mann-Whitney test with *p<0.05 **p<0.01, ***p<0.005 and ****p<0.001. D) Cells were treated overnight in 6 well plates, followed by lifting and staining with MitoSOX Red and flow cytometry using the FACs Fortessa (minimum 10,000 events acquired). Mean fluorescent intensity (MFI) is shown in inset. Where relevant, data was compared using a Mann-Whitney test with *p<0.05 **p<0.01, ***p<0.005 and ****p<0.001.

We next assessed whether activation of glycogen synthesis pathway contributed to the increased glycogenesis following CP91149 treatment. Glycogenesis is largely mediated by GS, which can be phosphorylated directly by activated glycogen synthase kinase 3B (GSK-3B), with phosphorylation serving to inactivate the enzyme and reduce synthesis (4,36). This axis can also be inhibited by insulin-driven activation of the PI3K/Akt pathway, which stimulates glycogen synthesis (4,36). Thus, glycogen synthesis is mediated by a balance of activating signals (i.e. Akt) and inactivating signals (i.e. GS and GSK). To assess the relative modulation of these signals with CP91149 treatment, immunoblotting was performed on KOC7C and 786-O cells treated for 24 hours with 20 μM or 100 μM CP91149, for phosphorylated GS, GSK, and AKT. Figure 2B illustrates that after 24 hours of treatment with CP91149, there is a marked increase in AKT activation at Ser^473^ for all three lines treated with 100 μM CP91149, suggesting a favouring of glycogen synthesis. Importantly, there is also a less-pronounced increase in the activation of GS (Ser^641^) with OVTOKO and KOC7C cells and GSK-3β(Ser^9^) with 786-O cells. This perhaps suggests that the inhibitor did not exert its maximal effect in 24 hours, or maybe indicates that signals for glycogenolysis still exist, thus highlighting the multi-faceted homeostatic control of the glycogen metabolism axis.

We then assessed the downstream effects of CP91149, by measuring ATP production in ccOC cells OVTOKO and KOC7C using the Seahorse ATP Rate Assay, which enables the delineation of ATP production from the routes of glycolysis and mitochondrial oxidative phosphorylation. The two ccOC lines were selected because they displayed the smallest increase in glycogen levels, leading us to postulate that the inhibitor was already exerting an effect downstream by 24 hours. Figure 2C shows that in ccOC lines OVTOKO and KOC7C, 100 μM CP91149 caused a reduction in ATP levels derived from both glycolysis and mitochondrial oxidative phosphorylation. Interestingly, mitochondrial oxidative phosphorylation was more acutely reduced by the inhibitor, which may be reflective of an increased dependence of these cells on the pathway, or a specific impact of the inhibitor on mitochondrial phenotypes such as apoptosis. In support of the latter hypothesis, both *pygl* silencing and CP91149 have previously been shown to specifically elicit mitochondrial ROS in glioma and breast cancer lines *in vitro* to elicit apoptotic cell death (7,32). This is thought to occur because PYG modulates glucose entry into the pentose phosphate pathway (PPP), which facilitates NADPH generation and subsequent ROS scavenging (7). To assess whether the inhibitor induced a similar phenotype in CCCs, the MitoSox superoxide sensor was employed, and ROS formation was assessed. Cells were treated for 24 hours with 20 μM or 100 μM CP91149, lifted, stained with the MitoSox dye, and then fluorescence measured flow cytometrically. As expected, CP91149 elicited a dose-dependent increase in MitoSox fluorescence after 24 hours of treatment, in both the ccOC and ccRCC lines, with a small increase in MFI with 20 μM, and a larger increase (∼double for OVTOKO, KOC7C and A498, ∼20% for 786-O), observed with 100 μM of the inhibitor (Figure 2D). This suggests that CP91149 is also modulating PPP flux to increase ROS in CCC lines.

### Low dose CP91149 specifically synergises with SOC OC chemotherapy paclitaxel in CCCs to elicit non-apoptotic cell death

Widespread chemoresistance remains one of the key prognostic hurdles for CCCs: for ccOC the platinum-taxol regime is largely ineffective, whereas ccRCC are highly resistant to first line chemo- and radio-therapies and are often refractory to TKI therapies (37,38). Because glycogen has been implicated in CCC chemoresistance in the past, we sought to assess whether PYG inhibition could synergise with existing standard of care therapies. A fixed dose of CP91149 was adopted – 20 μM – relatively ineffective as a single agent, alongside a dose range of kidney cancer therapy sunitinib and ovarian cancer chemotherapies carboplatin and paclitaxel. Cells were treated for 72 hours, followed by fixation, staining with DAPI, and automated cell counting. Synergism was measured using the Chou-Talalay Combination Index (CI) method, that calculates a CI based on median effects, where CI<1 indicates synergism, CI=1 indicates additive effects, and CI>1 indicates antagonistic effects (39). Figure 3A highlights the dose-response curves for ovarian cancer chemotherapy carboplatin alongside 20 μM CP91149 for ccOC lines OVTOKO and KOC7C, whilst Figure 3C highlights the combination of sunitinib with CP91149 in ccRCC lines 786-O and A498. CP91149 does not seem to synergise with either carboplatin nor sunitinib, as evidenced by the minimal shift in both dose-response curves and GI_50_ values. This is also reflected in the CI values in Figures 3B and 3D respectively, although there are weak suggestions of synergism (CI<0.9) with low-doses of carboplatin and sunitinib with CP91149, however these are likely false-positives that have been previously suggested to occur in low-order synergism cases using the CI method (40,41).

**Figure 3.**
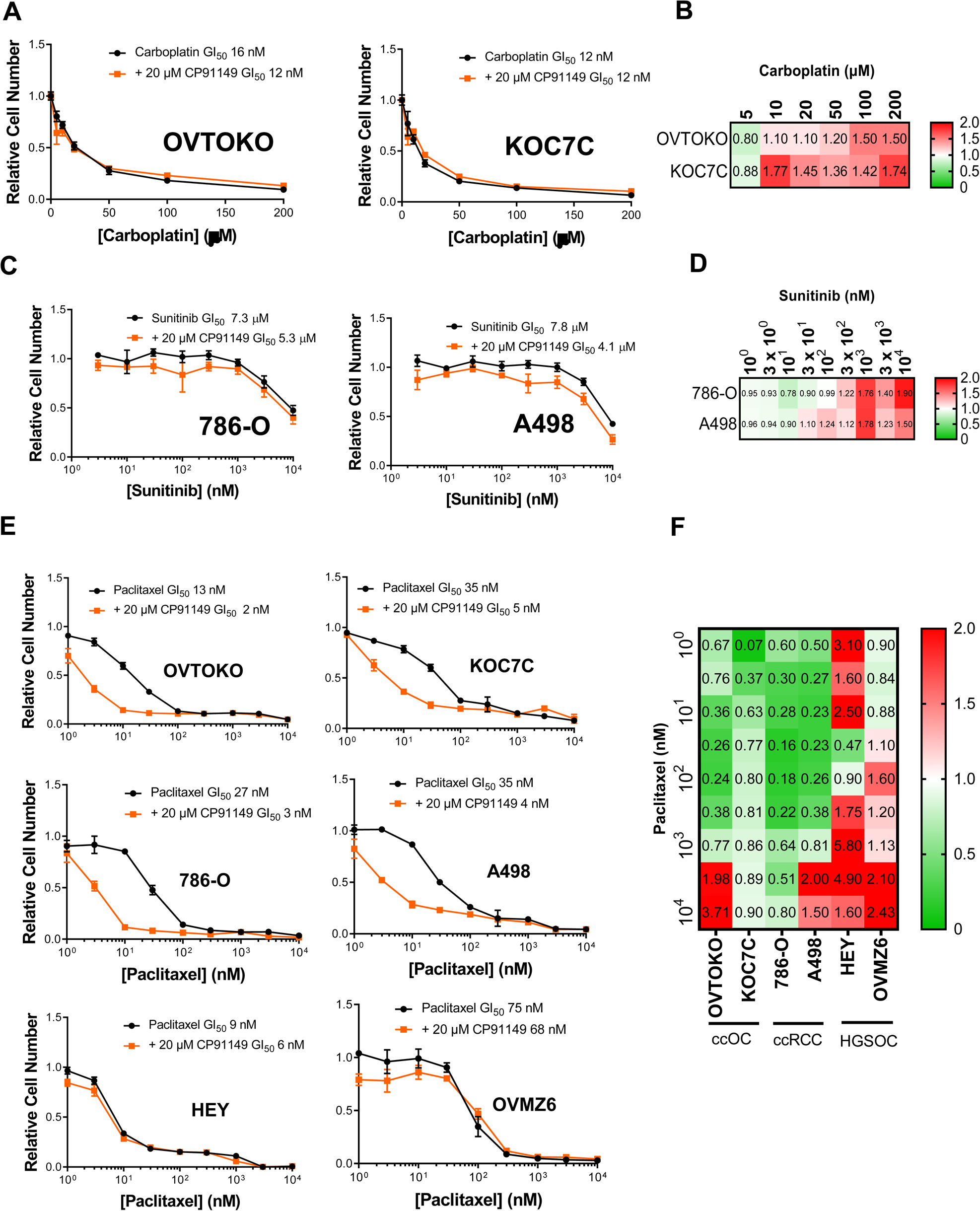
Low dose CP91149 specifically synergises with SOC OC chemotherapy paclitaxel in CCCs. **A)** Cells were treated with indicated concentrations of carboplatin in the presence or absence of 20 μM CP91149 for 72 hours, after which time they were fixed, stained with DAPI, imaged using the Incell 6500 system and analysed using CellProfiler software. B) Combination Index (CI) values were computed for cells treated with carboplatin alongside CP91149 using the Chou-Talalay Method. Data is presented in heatmap format with green indicating synergism and CI<1, white additive effects, and red antagonism. **C)** and **D)** As in A) and B) however cells were treated with varied doses of sunitinib alongside 20 μM CP91149. **E)** and **F)** As in A) and B) however cells were treated with varied doses of paclitaxel alongside 20 μM CP91149.

Contrastingly, we found that 20 μM CP91149 caused a marked downward shift in the paclitaxel dose-response in both ccOC lines OVTOKO and KOC7C and ccRCC cell lines 786-O and A498 (Figure 3E). This was also corroborated by the change in GI_50_ values for the lines: a 7-fold reduction for ccOC lines OVTOKO and KOC7C, and a 9-fold reduction for the ccRCC lines 786-O and A498. Specific synergism with paclitaxel was an unexpected finding, and hence we tested high grade serous ovarian cancer (HGSOC) lines HEY and OVMZ6 to assess whether this was a CCC-specific effect. Importantly, neither lines displayed the same marked shifts in dose-response curves nor GI_50_ and CI values, indicating that synergism between paclitaxel and CP91149 is a CCC-specific phenomenon. These trends were also reflected in the CI values, which suggested strong synergism in all cases but the two highest paclitaxel doses for OVTOKO and A498, which probably speaks to the monotherapy efficacy of paclitaxel at these concentrations (Figure 3F). in contrast, HEY and OVMZ6 cells did not display strong suggestions of synergism.

We also validated that this synergistic effect was specific to PYG by adopting a second PYG inhibitor, targeting a different site (Figure S1). We used BAY-R3401, which targets the AMP binding site of glycogen phosphorylase (42). We noted that BAY-R3401 similarly elicited a sharp drop in the dose-response curves of all four CCC lines (Figure S1A), eliciting a 20-fold reduction in GI_50_ for OVTOKO and KOC7C cells, a 10-fold reduction for 786-O cells, and a 5-fold reduction for A498 cells; a trend also reflected in the CI values for the combination (Figure S1B).

### Combination efficacy of paclitaxel and CP91149 is retained in ccOC cells stably resistant to paclitaxel

We next explored the efficacy of this drug combination using the MTT cell viability assay and clonogenic potential assay, to assess whether the potency of the combination was also reflected in different settings of cell growth. We employed GI_50_ values of paclitaxel for each line alongside 20 μM of CP91149. Viability of cells treated for 72 hours is shown in Figure 4A, which illustrates that the combination is more potent than either monotherapy for all 4 lines (p<0.05). Similarly, assessing clonogenic potential, the combination therapy elicited a near-complete ablation of colony-forming capacity of all four lines assessed (Figure 4B). Here, it is interesting to note that GI_50_ vales of paclitaxel alone had differing effects on the four lines: a minimal reduction in line OVTOKO (20%), more in KOC7C and 786-O (60%), and a marked effect for A498 (80%). This could be reflective of differing capacities to deal with the long-term effects of paclitaxel even after the drug’s withdrawal.

**Figure 4.**
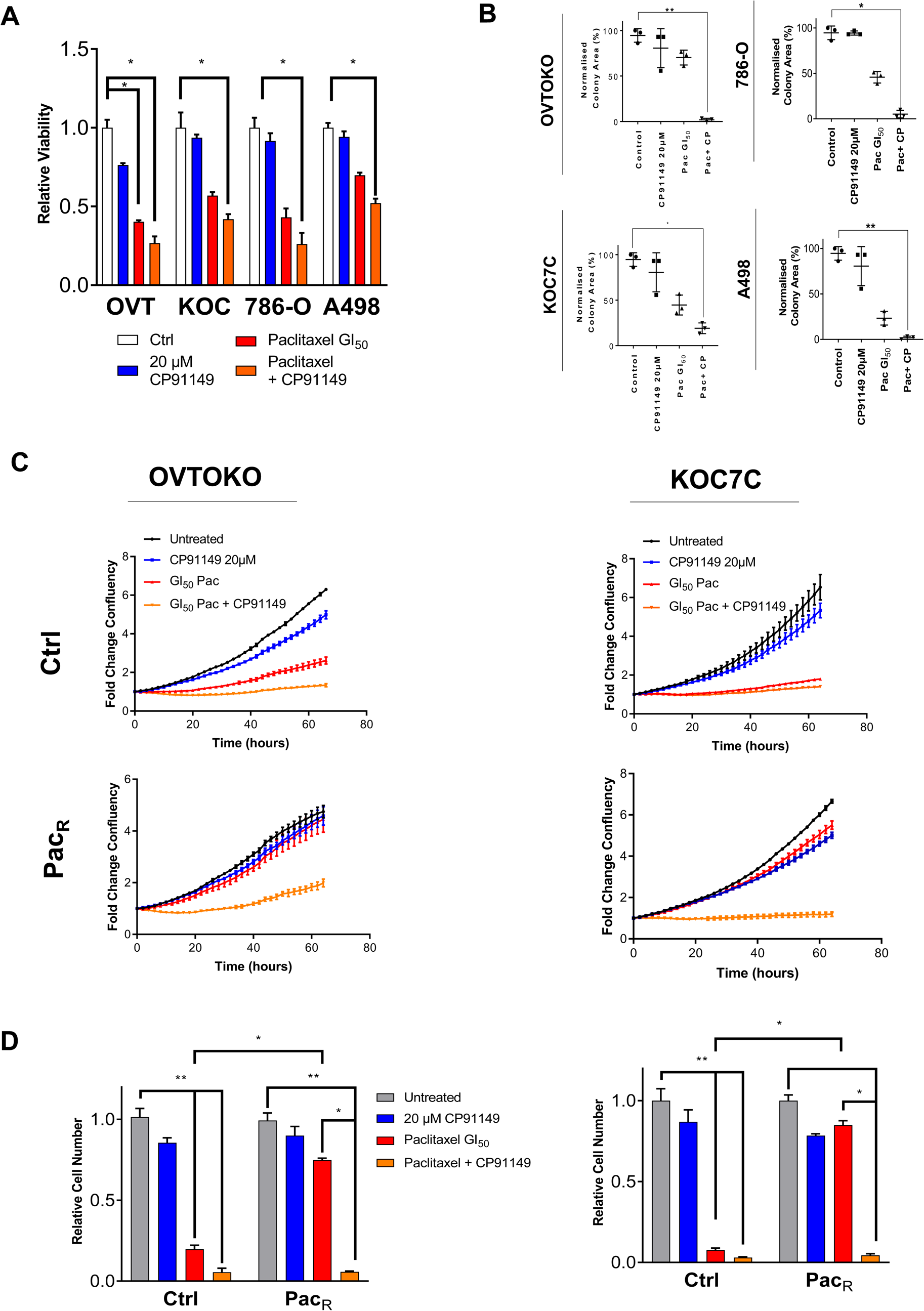
Combination efficacy of paclitaxel and CP91149 is retained in cell viability and colony formation assays, and in cells rendered stably resistant to paclitaxel. **A)** Cells were treated with indicated concentrations of paclitaxel in the presence or absence of 20 μM CP91149 for 72 hours, and viability was measured using the CellTiter AQueous One Solution MTT assay and normalized to untreated control. **B)** Cells were treated in 6 well plates with indicated drug doses for 72 hours, after which time they were lifted and re-seeded in drug-free media for 14 days. Colonies were fixed and stained with crystal violet, and colony area calculated using the ImageJ software. **C)** Paclitaxel resistant cells (Pac_R_) were generated by stepwise increase in paclitaxel concentration in culture between-drug free holiday periods to restore proliferative potential. Cells were considered stably resistant when they showed logarithmic growth in the presence of GI_50_ levels of paclitaxel. For assays, both Control and PacR cells were treated for 72 hours with indicated drug concentrations and imaged every two hours using the Incucyte S3 system. **D)** After 72 hours, cells were fixed, stained with DAPI and enumerated using the Incell 6500 system and CellProfiler software. Data is presented as mean ± SEM and was analysed using a Mann-Whitney test with *p<0.05 **p<0.01, ***p<0.005 and ****p<0.001.

Particularly for OC chemotherapy, poor prognosis is often characterised by primary disease response followed by disease relapse (43,44). Over subsequent chemotherapy doses, a resistant pool of drug-tolerant cells emerges, that are highly resistant to both the primary chemotherapy as well as other chemotherapy regimens (45). Given their importance to patient prognosis, we sought to generate paclitaxel resistant (Pac_R_) persister ccOC cells and assess the efficacy of the paclitaxel and CP91149 combination in these cells. OVTOKO and KOC7C cells were treated in culture with increasing doses of paclitaxel nestled between drug-free holiday periods, akin to what occurs in the clinical setting (45,46). Cells were deemed paclitaxel resistant when they displayed exponential growth in the presence of GI_50_ levels of the compound. Then, Control and Pac_R_ cells were treated with GI_50_ levels of paclitaxel alongside 20 μM CP91149. Change in confluency was measured using the Incucyte S3 system and endpoint cell number was also enumerated after 72 hours. Both Figure 4C and 4D show that OVTOKO and KOC7C Pac_R_ cells were robustly resistant to GI_50_ levels of paclitaxel. The paclitaxel response curves for Pac_R_ cells in Figure 4C revert nearly the same growth rate as control cells, whereas endpoint cell numbers highlight that the dose that caused 90% reduction in growth for control cells only reduced growth by 30% in Pac_R_ cells. Interestingly, assessing the combination of paclitaxel with CP91149, Pac_R_ cells were equally sensitive to control cells, illustrated both in terms of time-lapse confluence (Figure 4C) and endpoint cell number (Figure 4D). This highlights the potency of the drug combination, and potentially insinuates that PYG has a direct impact on paclitaxel resistance. Given that acquired chemoresistance is often a cause of poor prognosis for cancer patients, including ccOC and ccRCC patients, the observation that PYG inhibition can revert paclitaxel resistance can prove therapeutically useful (47,48).

### Combination-treated cells undergo ferroptotic cell death

Because of the magnitude of the effect for paclitaxel combined with CP91149, as well as the combination efficacy being retained in Pac_R_ cells, we wanted to further investigate the cause of this synergism. First, we assessed the mode of cell death elicited by combined treatment of paclitaxel and CP91149. Once again, time-lapse fluorescence microscopy was adopted using Annexin-V and PI. Cells (OVTOKO and KOC7C ccOC cells and 786-O and A498 ccRCC cells) were treated with GI_50_ concentrations of paclitaxel alongside 20 μM CP91149 and imaged every 30 minutes using the Incucyte S3 system as above. Cells were demarcated into AV+PI, AV-PI+, AV+PI+ or AV-PI-, and these were plotted with time; the frequencies of these populations for cells treated with paclitaxel and 20 μM CP91149 are shown in Figure 5A. Interestingly, in contrast to the early AV positivity followed by late dual positivity observed for CP91149-treated cells (Figure 1D), it was observed that for all four lines, cells treated with combination therapy underwent simultaneous annexin-V-PI dual-positivity (Figure 5A). This suggests that phosphatidylserine exposure does not preclude membrane rupture but rather occur alongside it, or even as a result of it. Such a trend is unlikely to be seen with apoptotic cell death, and is more likely reflective of non-apoptotic, necrotic modes of cells death (34,49–51).

**Figure 5.**
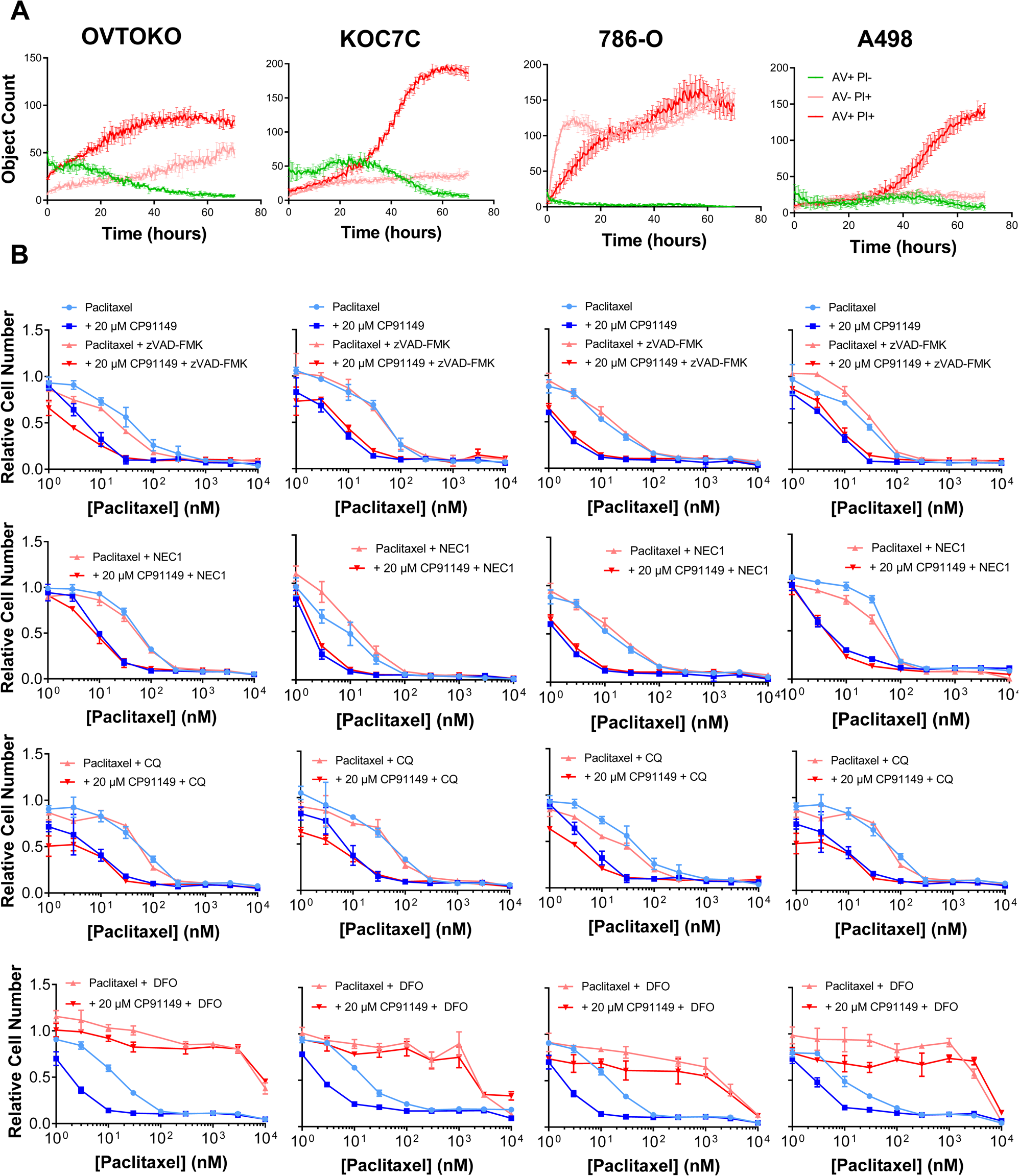
Paclitaxel and CP91149-treated cells potentially undergo ferroptotic cell death. **A)** Cells, pre-incubated with annexin V-FITC and propidium iodide, were treated with GI50 levels of paclitaxel alongside 20 μM CP91149 and imaged every two hours using the Incucyte S3 system. Single cells were analysed using Incucyte software and defined as either AV+PI-, AV-PI+, AV+PI+ or AV-PI-, and cell populations are plotted with time. **B)** Cells were treated for indicated concentrations of paclitaxel in the presence or absence of CP91149 (20 μM), necrostatin-1 (NEC-1; 1 μM), deferoxamine (DFO; 100 μM), zVAD-FMK (20 μM) or chloroquine (CQ; 50 μM) for 72 hours, after which time they were fixed, stained with DAPI, and enumerated using the Incell 6500 system and associated CellProfiler software.

We sought to determine the mode of non-apoptotic cell death elicited by treatment with paclitaxel and CP91149. We attempted to rescue combination efficacy with inhibitors for several cell death pathways: chloroquine for autophagic cell death, necrostatin-1 for necroptotic cell death, deferoxamine for ferroptotic cell death, and z-VAD FMK for apoptotic cell death (Figure 5B). We reasoned that a rescuing of the observed synergism would be suggestive of the cell death pathway of combination-treated cells. Cells were treated with the combination of paclitaxel and CP91149 in the presence of rescue agents, and endpoint cell number was calculated as previously. Figure 5B shows that neither zVAD-FMK, necrostatin-1 nor chloroquine elicited any marked shift in the dose-response curves, suggesting that cells are not undergoing apoptotic, necroptotic, or autophagic cell death. Notably, iron chelator deferoxamine was the only agent to consistently reduce combination efficacy, as demonstrated by an upward shift of both the paclitaxel-treated and combination-treated dose-response curves in all four cell lines (Figure 5B), suggesting cells were undergoing ferroptotic death. Ferroptosis is an iron-dependant form of cell death that culminates in peroxidation of the lipid membrane and is mediated by the glutathione peroxidase 4-driven antioxidant state of cells (52,53). Importantly, CCCs have previously been shown hypersensitive to ferroptosis, owing to a GPX4-dependant cell state (12). Furthermore, ferroptosis has previously been shown to be an achilles’ heel of drug-tolerant persister cells (45), which perhaps explains the efficacy of combination treatment in paclitaxel-resistant lines in Figure 4.

### CCC cells treated with combined CP91149 and paclitaxel display hallmarks of ferroptosis

We examined whether cells treated with paclitaxel and CP91149 were undergoing ferroptosis by assessing several markers that characterise the phenotype: mitochondrial hyperpolarisation, transferrin receptor translocation, increased labile iron, and lipid peroxidation. Mitochondrial hyperpolarisation has been shown to drive ferroptosis by facilitating the rapid depletion of the cellular antioxidant system (54). We examined mitochondrial membrane potential by fluorescence microscopy using the rhodamine dye TMRE. Cells were pre-incubated with TMRE for 30 minutes, treated with indicated compounds, and the steady-state TMRE fluorescence per cell was measured using the Incucyte S3. Figure 6A shows that there was a slight increase in TMRE fluorescence per cell for treatment with 20 μM of CP91149, and a robust increase for both paclitaxel and combination-treated cells. For OVTOKO this corresponded to a two-fold increase, for KOC7C and A498 a 5-fold increase, and for 786-O a 7-fold increase in TMRE fluorescence per cell. Hence, Figure 6A suggests that CP91149 and paclitaxel treatment increases mitochondrial membrane potential and corroborates ferroptosis.

**Figure 6.**
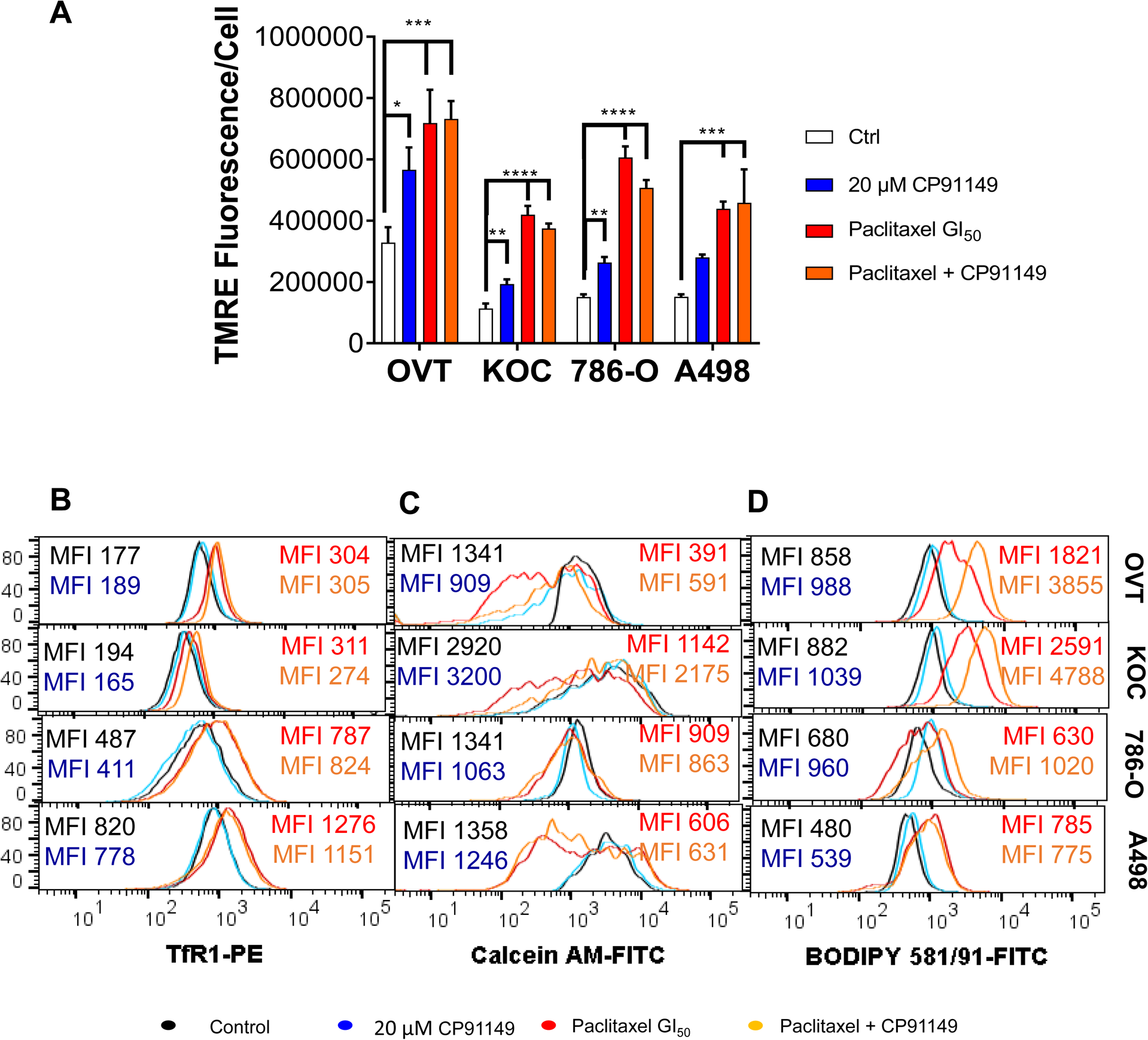
Paclitaxel and CP91149-treated cells display several hallmarks of ferroptosis. **A)** Cells were preincubated with 100 nM TMRE for 30 mins, after which time the relevant agents were added and cells imaged using the Incucyte S3 system. TMRE fluorescence/cell was calculated at 30 minutes post-treatment. **B)** Cells were treated and lifted as in B), this time blocked (30 mins 0.5% BSA in PBS) and incubated with anti-Cd71 TfR1 antibody (Biolegend) and at least 10,000 events acquired flow cytometrically using the FACS Fortessa system. **C)** Cells were treated for indicated drugs for 24 hours, after which time they were lifted, stained for 30 minutes with calcein-AM (100nM) and at least 10,000 events acquired flow cytometrically using the FACS Fortessa system**. D)** Cells were treated as in B), however here they were incubated with BODIPY 581/91 C11 lipid peroxidation sensor for 30 minutes, followed by lifting and at least 10,000 events acquired flow cytometrically using the FACS Fortessa system. Where relevant, data is presented as mean ± SEM and was analysed using a Mann-Whitney test with *p<0.05 **p<0.01, ***p<0.005 and ****p<0.001.

Transferrin receptor membrane translocation and the labile iron pool facilitate iron entry into cells and iron-catalysed lipid peroxidation and are thus also markers for ferroptosis (53,55). Both were measured by flow cytometry on cells treated with paclitaxel and CP91149 for 24 hours. Membrane transferrin receptor (TfR1) has recently been shown to be a faithful marker for ferroptosis induction, presumably because transferrin facilitates iron entry and thus facilitates increases in the labile iron pool (55). We measured membrane expression of TfR1 using anti-human CD71-PE conjugate by flow cytometry (Figure 6B). We found that both paclitaxel and the combination treatment resulted in increased TfR1 expression for all four lines by approximately 40%. This suggests an increase in labile iron and a priming for ferroptosis.

We next measured labile iron pool using the dye calcein-AM (Figure 6C) which is robustly quenched by intracellular iron (52). Figure 6C shows that calcein fluorescence decreased with both paclitaxel and combination treatment for all four lines, quite robustly for ccOC lines OVTOKO and KOC7C and ccRCC line A498 (∼60% reduction), and slightly for ccRCC line 786-O (∼25%). This corroborates trends for TfR1 translocation and suggests paclitaxel and combination treatment for all four lines facilitates ferroptosis.

The final cytotoxic event of ferroptosis is peroxidation of the lipid membrane, which disrupts membrane integrity and thus elicits cell death (52). Lipid peroxidation can also be measured flow cytometrically, with the BODIPY dye 581/91-C11, a specific marker of peroxidised lipids. BODIPY-C11 fluorescence was measured in cells treated for 24 hours with 20 μM of CP91149, GI_50_ paclitaxel, or a combination. Figure 6D shows that paclitaxel caused a slight shift in BODIPY-FITC fluorescence for OVTOKO, KOC7C, and A498 cells, and no foreseeable shift for 786-O cells. This is reflected in a doubling of MFI values for OVTOKO and A498, and a 3-fold increase for KOC7C. Combination therapy elicited a greater shift for OVTOKO, KOC7C and 786-O cells, while it elicited an equivalent shift to paclitaxel in A498 cells. This is reflected in a doubling of MFI values for 786-O and A498 cells, and a 4-fold increase for OVTOKO and KOC7C cells compared to control. This increased lipid peroxidation supports the trajectory of ferroptotic cell death.

Our data suggests cotreatment with paclitaxel and CP91149 causes ferroptosis, however it remains to be seen how this drug combination is causing ferroptotic cell death in specifically CCCs. Interestingly, paclitaxel monotherapy augurs many of the hallmarks of ferroptosis, including increased mitochondrial membrane potential, increased labile iron and increased membrane TfR1 expression (Figure 6). The ability of taxol to increase mitochondrial membrane potential has been reported previously and suggested to be the result of a decrease in cytoplasmic free tubulin, which can act to depolarise mitochondria (56). It is possible that this this presents a sufficient ferroptotic stimulus to CCCs, which are already hypersensitive to ferroptosis, when PYG is inhibited to a sufficient extent to modulate redox potential. If this were the case, one would hypothesise that different tubulin- and cell cycle-targeted drugs would not engender similar synergism.

To test this hypothesis, we tried both the microtubule destabiliser nocodazole and the PLK-1 inhibitor BI2596 in combination with 20μM CP91149. Nocodazole acts in the opposite way to paclitaxel, destabilising microtubules and thus increasing free tubulin (56). BI2596 inhibits the enzyme polo-like kinase 1 (PLK1), which serves as an early trigger to G2/M transition, and thus inhibits mitosis via a different mechanism altogether to paclitaxel and nocodazole (57). We treated OVTOKO and KOC7C ccOC cells with both inhibitors alongside CP91149 (Supplementary Figure 2). Supplementary Figure 2 shows that the curves of both nocodazole and BI2536 shifted downwards with the addition of CP91149, but not to the same marked extent as paclitaxel. For nocodazole, the addition of CP91149 elicited a small shift in GI_50_ values (20% reduction); whilst for BI2536 it was a more pronounced effect – 60% for paclitaxel and 70% for KOC7C. The observation that nocodazole – which acts similarly to paclitaxel – does not similarly synergise with CP91149 is important, as it suggests synergism is specific to agents that decrease free tubulin. However, it is interesting to note that the dose-response curve of PLK1 inhibitor BI2536 did shift more markedly with CP91149, which may indicate additional mechanisms of synergy between PYG inhibitors and cell-cycle targeted drugs. This would support suggestions of the importance of PYG in cell cycle progression in both cancerous and normal physiology settings (3,7,58).

## DISCUSSION

The present work investigated CP91149, an inhibitor of the indole carboxamide site of all PYG isoforms. The inhibitor as a monotherapy induced dose dependant effects in CCC lines akin to that which had been previously shown in non CCC lines, specifically hepatocellular carcinoma and PDAC lines (30,32). Interestingly however, synergism with microtubule disrupting chemotherapy paclitaxel was noted only in CCC lines. Given that small-molecule inhibitors can elicit off-target effects, we verified our synergistic efficacy with a second inhibitor for a different PYG site; BAY R3401, an allosteric site inhibitor of the phosphorylase-α subunit. A necessary future direction for the present work is to verify this synergistic efficacy using genetic approaches to PYG silencing, however such approaches may be encumbered by compensation between isoforms as well as the potential that different phenotypic effects may be elicited by inhibiting the enzyme constitutively versus at a specific point in time.

The combination of paclitaxel with PYG inhibition seems to promote ferroptotic cell death, given that cytotoxicity was rescued by the iron chelator deferoxamine, and that combination therapy displayed many of the signatures of ferroptosis. It is not entirely clear how this combination therapy is eliciting ferroptosis. That the combination of paclitaxel with CP91149 is specifically potent in CCCs is perhaps explained by the hypersensitivity of the histotype to ferroptosis, owing to a pseudohypoxia-based reliance on glutathione peroxidase 4 (GPX4), a master gatekeeper of ferroptosis (12,59). Zou and colleagues (2019) recently found that HIFs selectively enrich polyunsaturated lipids, the rate-limiting substrates for lipid peroxidation, in CCCs, which render cells hypersensitive to the induction of ferroptosis by the inhibition of GPX4. In the present setting, we hypothesise that PYG inhibition lends itself to ferroptosis by hampering the antioxidant capacity of CCCs, a hypothesis supported by a CP91149-based increase in mitochondrial reactive oxygen species. Interestingly, glycogen metabolism has been implicated in ferroptosis in the past, with an upstream modulator of PYG, phosphorylase kinase G (PHKG2), recently implicated as a modulator of labile iron and ferroptosis by Yang and colleagues (2016) (52). Importantly though, the authors noted a ferroptosis-rescuing effect with PHKG2 silencing and postulated this to be due to a moonlighting effect of the enzyme, separate to its role in glycogenolysis, given that this rescue was not recapitulated by PYG inhibition (52). Importantly, the authors did note a ferroptosis sensitising effect with CP91149, a phenomenon that supports the findings of the present work (52).

The role of paclitaxel in eliciting ferroptosis is an intriguing one; although taxol has previously been shown to cause ferroptosis (53), the link between cell cycle arrest and ferroptosis is currently tenuous. Moreover, in the present work we have shown that two other cell cycle inhibitors – microtubule destabiliser nocodazole and PLK1 inhibitor BI-2596 – do not similarly synergise with PYG inhibition nor seem to augur ferroptotic cell death. We hypothesise that the specificity of paclitaxel may be due to the drug’s unique effect on free tubulin; as a microtubule stabiliser paclitaxel diminishes the amount of free tubulin the cytosol (56). Free tubulin can act directly on mitochondrial voltage-dependant anion channels to depolarise mitochondria (56). A lack of free tubulin can hyperpolarise mitochondria, which has been shown to sensitise cells to ferroptosis (54). In this regard, it may also be useful to assess whether paclitaxel synergises with existing ferroptosis inducers, such as those targeting system XC or GPX4. These compounds have thus far been encumbered pre-clinically by poor bioavailability and pharmacokinetic properties (59–61); the combining them with existing chemotherapeutic agents may potentially provide a therapeutic window.

## CONCLUSIONS

In conclusion, the present work shows that PYG inhibition may be improve the efficacy of paclitaxel chemotherapy against CCCs. However future work is required to decide and assess the ideal dosing regimen for *in vivo* studies.

## ABBREVIATIONS

CCC: Clear cell carcinoma
ccOC: Clear cell ovarian cancer
ccRCC: Clear cell renal cell carcinoma
GPX4: glutathione peroxidase
GS: glycogen synthase
PYG: Glycogen Phosphorylase
CI: Combination Index
HGSOC: High Grade Serous Ovarian Cancer
PacR: Paclitaxel Resistant

---

**Figure S1.**
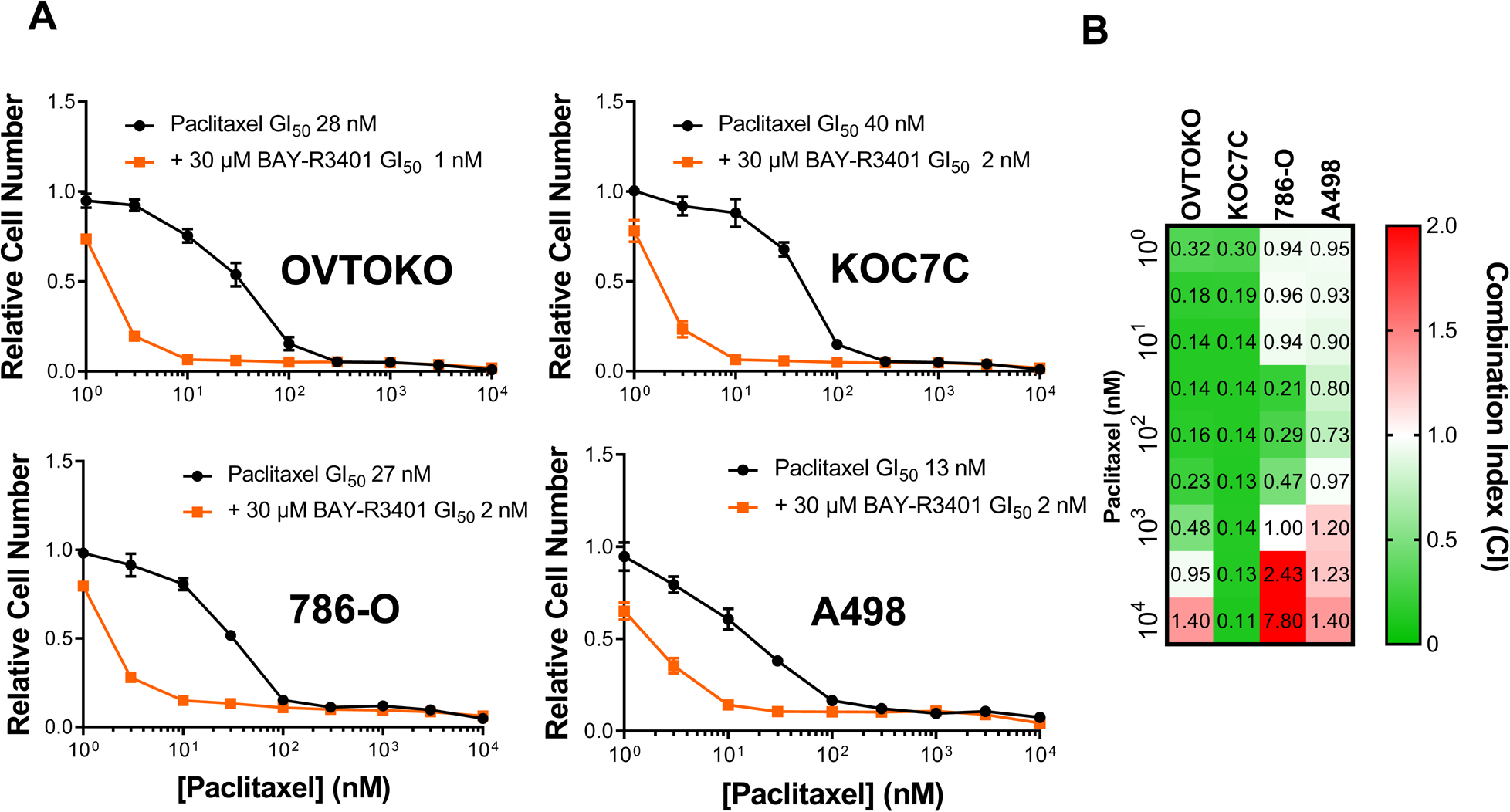

**Figure S2.**
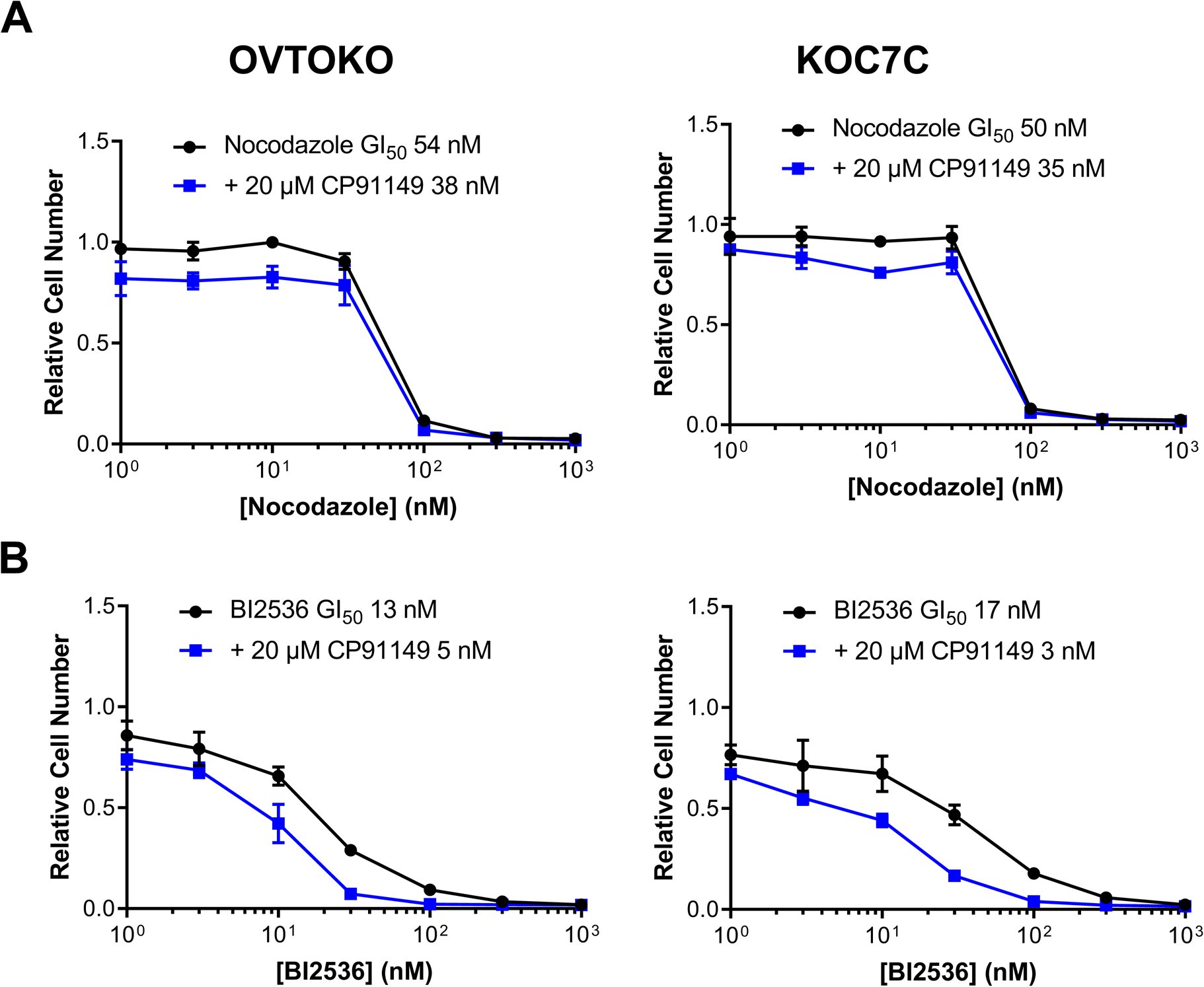

